# Resilience to oxidative and nitrosative stress is mediated by the stressosome, RsbP and SigB in *Bacillus subtilis.*

**DOI:** 10.1101/460303

**Authors:** Vina Tran, Kara Geraci, Giovanni Midili, William Satterwhite, Rachel Wright, Carla Yaneth Bonilla

**Author notes:** Corresponding author: Carla Y. Bonilla, Assistant Professor, Biology Department, Gonzaga University, 502 E Boone Avenue, Spokane WA, 99253.

## Abstract

A bacterium’s ability to thrive in the presence of multiple environmental stressors simultaneously determines its resilience. We showed that activation of the SigB-controlled general stress response by mild environmental or nutritional stress provided significant cross-protection to subsequent lethal oxidative, disulfide and nitrosative stress exposure. SigB activation is mediated via the stressosome and RsbP, the main conduits of environmental and nutritional stress, respectively. Cells exposed to mild environmental stress while lacking the major stressosome components RsbT or RsbRA were highly sensitive to subsequent oxidative stress, whereas *rsbRB, rsbRC, rsbRD* and *ytvA* null mutants showed a spectrum of sensitivity, confirming their redundant roles and suggesting they could modulate the signal generated by environmental stress or oxidative stress. Furthermore, from mutant analysis we infer that RsbRA phosphorylation by RsbT was important for this cross-resistance to oxidative stress. By contrast, cells encountering stationary phase stress required RsbP but not RsbT to survive subsequent oxidative stress caused by hydrogen peroxide and diamide. Interestingly, optimum cross-protection against nitrosative stress caused by SNP required SigB but not the known regulators, RsbT and RsbP, suggesting an additional and as yet uncharacterized route of SigB activation independent of the known environmental and energy-stress pathways. Together, these results provide a mechanism for how *Bacillus subtilis* promotes enhanced resistance against lethal oxidative stress during likely physiologically relevant conditions such as mild environmental or nutrient stress.

## Importance

The *Bacillus subtilis* general stress response is a model for gram-positive pathogens because the regulators are conserved, and the Sigma factor, SigB, controls expression of virulence genes in *Listeria monocytogenes.* We showed that *B. subtilis* SigB promotes survival to oxidative, disulfide and nitrosative stress through priming or cross-protection. Moreover, when cells were exposed to nitrosative stress, priming was SigB dependent, yet the known regulators of SigB were not required, suggesting an alternative mode of SigB activation during nitrosative stress. Importantly, we showed the first genetic requirements of stressosome genes, *rsbRB* and *rsbRD,* during oxidative stress cross-protection not explained by environmental stress activation, suggesting a role for stressosome proteins during oxidative stress and advancing the role of SigB during antioxidant protection.

## Introduction

The resilience of bacteria to environmental stressors allows them to survive during constantly changing conditions (1). Resilience comes about due to cross-protection or priming, which is when bacteria face mild stress that prepares them for future lethal stress, whether or not the stresses are related. Microbes commonly use this phenomenon in order to survive their dynamic environments (2). Priming is especially beneficial for pathogens because it increases their fitness in the face of the host immune system, which deploys an oxidative burst meant to kill the pathogen (1). Changes to gene expression induced by stress are important for the acquired cross-protection that will let those preprogrammed cells thrive in the presence of a further lethal stressor. Most environmental stresses, such as osmotic shock and temperature shifts, can cause priming and have the highest protective effect against oxidative stress (2). Therefore, evolutionarily conserved mechanisms of bacterial survival, environmental stress induced gene expression and the proteins involved in these processes could have broader biological significance than previously recognized.

*Bacillus subtilis* uses multiple sigma factors to cope with the changes to its surroundings (3). Some sigma factors are dedicated to specific stressors, but the general stress sigma factor, SigB is responsible for the adaptation to the widest type of environmental conditions and therefore is an important protein during priming (4). Environmental conditions include high and low temperature, alkaline and acidic environments, osmotic stress and changes in carbon sources, ATP levels and oxidative stress (5-8). The activation of SigB leads to the differential expression of 196 genes with diverse biochemical functions giving cells resistance to multiple stresses, an important aspect of priming (9). Moreover, survival is enhanced in the presence of reactive oxygen species (ROS) when priming is triggered by ethanol stress (10). In this comprehensive analysis of 94 individual SigB-dependent genes, Reder and colleagues showed priming protection to lethal levels of hydrogen peroxide. Cells carrying mutations in individual genes were first given non-lethal ethanol exposure, to trigger priming, followed by lethal levels of hydrogen peroxide and stress-induced tolerance was dependent on many SigB targets (10). It has also been shown that in the presence of oxidative stress alone, caused by hydrogen peroxide or Sodium Nitropruside (SNP), genes belonging to the SigB regulon are induced (11-14), suggesting that SigB is activated by the presence of oxidative stress signals. Furthermore, the need for SigB in resistance against oxidative stress is apparent in stationary phase cells, where exposure to hydrogen peroxide made *sigB* null cells more sensitive than wild type (15).However, the upstream mechanisms controlling SigB-dependent priming during ethanol exposure and in nutritionally stressed cells have not been addressed.

SigB activity is controlled by two pathways (Figure 1A), which independently sense nutritional and environmental stresses (16). Nutritional stress such as low ATP levels requires the activity of the hydrolase RsbQ and phosphatase RsbP, although the specific nutritional signal is unknown (17-19). Environmental stress uses the stressosome complex consisting of related, putative sensor proteins RsbRA, RsbRB, RsbRC, RsbRD, and YtvA, and the kinase RsbT and antagonist RsbS (20-23). The RsbR paralogs are candidate sensor proteins due to their amino acid sequence similarity to known sensing domains and their position on the 3D structure of the stressosome. The N’ termini, containing a non-heme globin domain are found externally in the structure while the STAS C’ terminal domains interact with RsbS and RsbT, potentially transmitting the environmental signal (24-26). Once the stress is sensed, such as ethanol or osmotic stress, RsbT is activated, phosphorylates RsbRA and RsbRB, and leaves the stressosome complex (20, 23, 27). The specific signal that initiates the signaling cascade remains unknown, but could be transmitted from the environment to the stressosome through the N’ termini of the RsbR proteins. Once released from the stressosome, RsbT activates the phosphatase RsbU through their direct interaction (28). Active RsbU dephosphorylates RsbV, promoting the partner switching of RsbW bound to SigB to the anti-sigma factor RsbV (29). SigB is normally associated with RsbW, but the dephosphorylation of RsbV causes RsbW to switch partners releasing SigB (30, 31). Nutritional stress works similarly promoting RsbV activation. Upon ATP level depletion the RsbP/RsbQ dimer becomes activated promoting RsbP phosphatase activity towards RsbV resulting in the activation of SigB by releasing RsbW (32). Once SigB is activated, at least 196 genes become differentially expressed leading to the production of important proteins that protect the cell in these stressful circumstances. Although their relative contribution to survival under environmental stresses including oxidative stress has been measured (6, 10), the role of individual stressosome components during extreme oxidative stress, cross-protection has not been determined.

**Figure 1.**
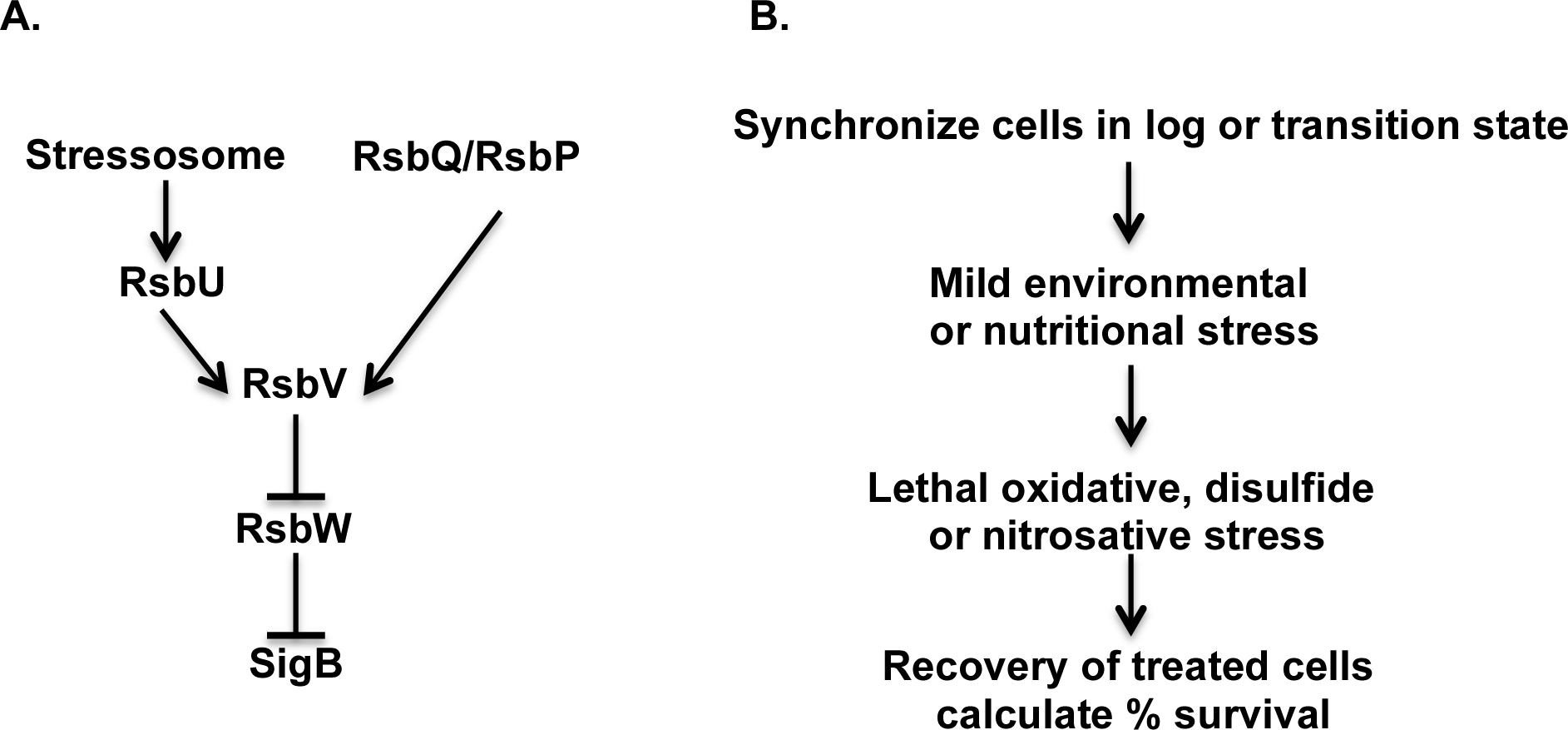
SigB activation pathway. Environmental stress and nutritional stress activate the stressosome and RsbP respectively. The phosphatase RsbU and RsbP activate the anti-anti sigma factor RsbV which then inhibits the anti-sigma factor RsbW, releasing SigB. (B) Experimental approach to test bacterial cross-protection to oxidative stress in log or transition state.

Here, measuring the ability of cells mutant in SigB-regulatory proteins to survive exposure to lethal Reactive Oxygen Species (ROS) and Reactive Nitrogen Species (RNS), we probed the role of key SigB regulators in promoting cross-protection during logarithmic and stationary phase. We showed that when priming is prevented by deleting the transducers of environmental and nutritional stress, cells became sensitive when placed in the presence of oxidative and nitrosative stress. In the case of nitrosative stress caused by SNP, *sigB* mutants were the most sensitive followed by individual and double *rsbT, rsbP* mutants. This result demonstrated the presence of SigB-dependent pathways responsible for stress protection seen in cells during nitrosative stress that are independent of the stressosome or the nutritional stress sensors. Moreover, we showed for the first time the effect of deleting individual stressosome genes in the physiology of priming suggesting that improper environmental stress signaling is detrimental to cells when dealing with extreme oxidative stress.

## Materials and Methods

*Bacterial strain construction.* Strains used in these experiments were either made as described in Table1, donated by the Bacillus Genetics Stock Center or courtesy of Dr. Chester Price at the University of California, Davis. Deletions of *rsbRA, rsbT, rsbP* were made by gene replacement with the Chloramphenicol or Kanamycin resistance cassettes from plasmid pGK67. PCR products were made containing 1000 base pairs of homologous regions upstream and downstream of each gene flanking the desired antibiotic resistance gene using NEB Q5 Polymerase. These PCR products were used to transform wild type cells, then antibiotic resistant transformants were confirmed by PCR of the desired mutation at the endogenous locus. In the case of the *rsbRA* deletion, reverse transcription PCR was performed to confirm that the insertion-deletion was not polar on the operon and that the *rsbT* and *rsbS* transcripts were still expressed. All other deletions were made by chromosomal transformation with DNA from strain PB804 containing the desired mutations of stressosome genes. Strain PB804 containing antibiotic marked deletions of *rsbR* genes was used to delete individual stressosome components and selected for single mutations. These strains were also confirmed by PCR of the endogenous locus of each gene. DNA isolation and plasmid preparations were performed using Zymo Reseach kits.

**Table 1.**
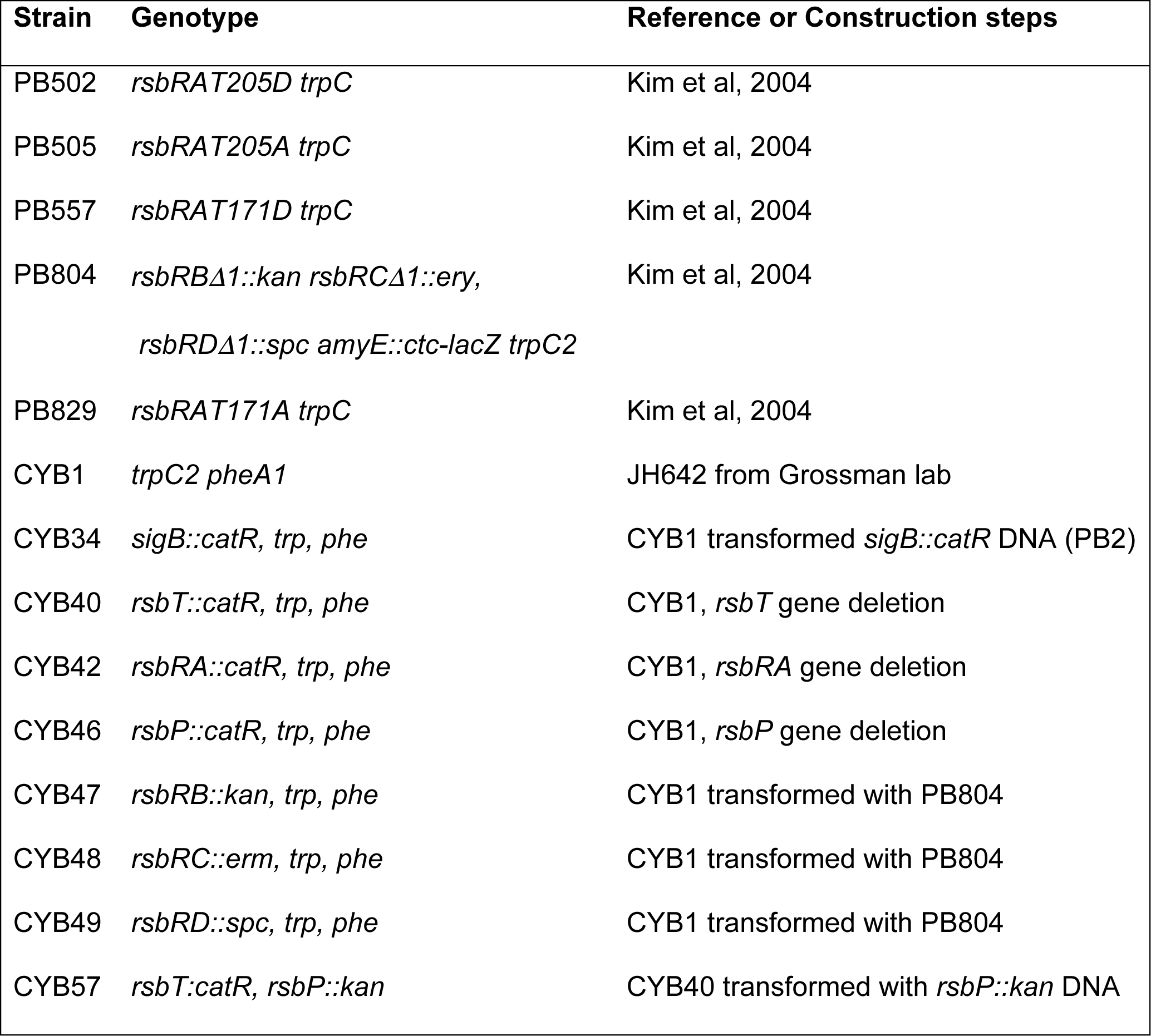
Strains used in experiments.

### Oxidative stress viability and calculations

*Bacillus subtilis* strains were grown in Basal Limitation Media (BLM) for experiments in exponential phase, or Glucose Limitation Media (GLM) for stationary phase treatment as previously described (33). BLM consists of 50 mM Tris, 15 mM(NH_4_)_2_SO_4_, 8 mM MgSO_4_, 27 mM KCl, and 7 mM sodium citrate (pH 7.5), 2 mM CaCl_2_, 1uM FeSO_4_, 10 uM MnSO_4_, 4.5 mM Potassium Glutamate, 0.1% glucose (or 0.05% for GLM), 0.6 mM KH_2_PO_4_ and 160 ug ml^-1^ each of Trp and Phe. Overnight cultures of *B. subtilis* were used to create starting cultures at OD_600_ readings of 0.05, incubated at 37°C while shaking at 300rpm until OD_600_ reached mid-log (∼0.4). At mid-log cultures were treated with 2% ethanol for 20 minutes, while shaking. These cultures were split and treated with either 5 mM H2O2, 45 mM diamide or 74 mM SNP. After 60 minutes, the cultures were serially diluted and plated on LB agar to recover at 37°C for 16 hours before colonies were counted. For oxidative stress induction during nutritional stress, cells were grown in GLM for the entire experiment and growth was monitored until cells reached transition state. One hour into transition state, 10 mM H_2_O_2_, 45 mM diamide, or 74 mM SNP was added for an additional 60 minutes while shaking. Both treated and untreated cultures were diluted and plated in LB agar plates and allowed to recover for 16 hours at 37°C. For each individual experiment, treated and untreated bacterial cultures were plated in triplicate, counted and averaged. To calculate percent survival, the number of colonies forming units under stress was divided by the number of colonies forming units without stress. The data shown represent percent viability means of a minimum of three experiments with standard error bars. To calculate the significance of the difference between the percent viability means of different strains, the data were subjected to Ordinary One-Way ANOVAs and all showed P values of 0.005 or lower. Tukey’s or Dunnett’s multiple comparison tests were performed to compare viabilities between strains. All strains were compared to wild type and to *sigB* nulls when appropriate.

## Results

### RsbT and RsbP are required for cross-protection to lethal reactive oxygen species

We set out to test the role of key SigB regulators, the stressosome and RsbP, during cross-protection to oxidative stress. Each pathway operates during different growth phases; logarithmic cells are sensitive to environmental stress, transmitted via the stressosome, and early stationary phase cells are nutritionally starved, a condition signaled via RsbP. We hypothesized that each regulator would be required to promote oxidative stress cross-protection in their respective growth phases; therefore we performed experiments in both log phase and early stationary phase using BLM and GLM respectively (Figure 1B). In order to test priming or cross-protection, in log phase, cells were primed with sub-lethal levels of ethanol and then treated with lethal hydrogen peroxide levels as previously shown (10). Wild type cells preadapted with ethanol were more resistant than cells that received hydrogen peroxide alone and their survival was dependent on SigB since *sigB* deleted cells were extremely sensitive (20 fold, decrease in surival) to the subsequent exposure to hydrogen peroxide (Figure 2A). Since resistance to oxidative stress in nutritionally stress cells was shown to depend on the alternative sigma factor SigB (15), we set out to identify the signaling pathway involved in the cross-protection in this phase. In nutritionally starved cell, RsbP/RsbQ are responsible for SigB activation, therefore we treated *rsbP* deleted cells with lethal amounts of hydrogen peroxide. *rsbP* mutants were highly sensitive to oxidative stress, similarly to *sigB* deleted cells (Figure 2B). In contrast, *rsbT* knock out cells were not sensitive and exhibited survival indistinguishable from wild type cells, demonstrating that the stressosome does not play a role in the stationary phase-induced oxidative stress cross-protection (Figure 2B).

**Figure 2.**
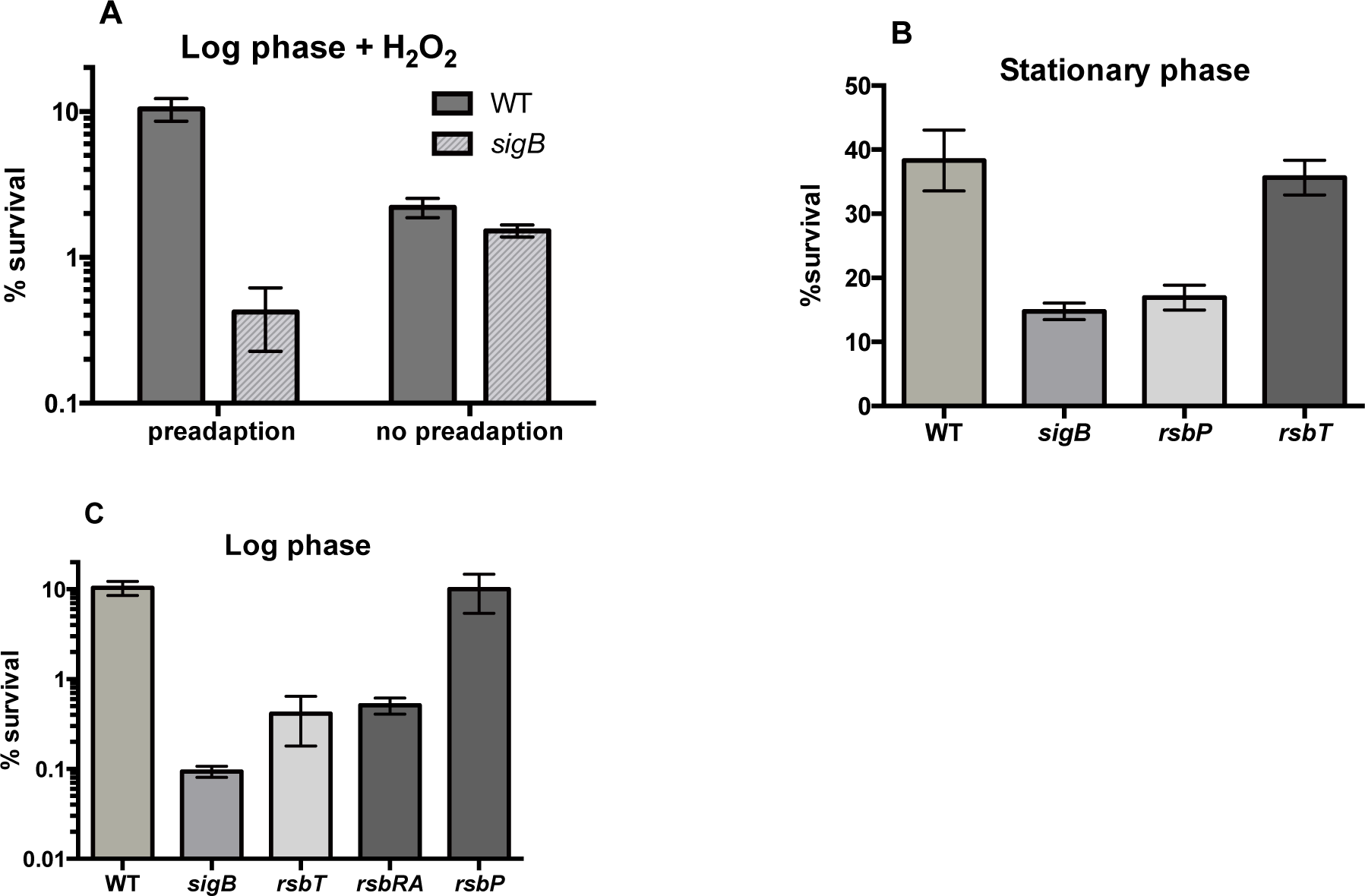
Environmental and nutritional stress protects cells from oxidative stress. (A) Cells were grown in BLM until OD_600nm_ 0.4 and were either preadapted with 2% ethanol for 20 minutes or received no preadaptation. Hydrogen peroxide was added at 5 mM final concentration for 1 hour before cells were allowed to recover overnight on LB plates. (B) Cells were grown in Glucose Limitation Media and monitored until transition state. One hour into transition state, cell were given 10 mM hydrogen peroxide for one hour and then plated. (C) Cells were grown to midlog in BLM, treated with 2% ethanol for 20 minutes and then given 5 mM hydrogen peroxide for one hour. Every experiment was done a minimum of three times and averaged; standard error bars are shown for all experiments. One-way ANOVAs were performed, followed by Tukey’s multiple comparison tests to determine statistical significant differences between means.

In order to test the role of environmental stress-activated SigB during priming against oxidative stress, we used mutations in members of the stressosome to assess their role in cross-protection. The kinase RsbT and the co-antagonist RsbRA were deleted individually and logarithmically growing cells were preadapted with mild ethanol stress before being given lethal levels of hydrogen peroxide. *rsbT* and *rsbRA* mutant cells (Figure 2C) were more sensitive than wild type cells and similarly sensitive to *sigB* deleted cells (ANOVA P value 0.0016, Tukey’s test showed no significant difference amongst *rsbT, rsbRA* and *sigB* cells) showing that the stressosome is important for the cross-protection that renders the cells resistant to oxidative stress. Deletion of *rsbP* had no effect on survival to hydrogen peroxide exposure (Figure 2C) demonstrating that RsbP is not required for cross-protection to oxidative stress in logarithmically growing cells likely due to not being activated by this stress.

### Stressosome components play different roles in the resilience to reactive oxygen species

The role of the stressosome in ROS cross-protection has never been tested, so we characterized mutants in individual stressosome components. The stressosome is made up of five paralog proteins RsbRA, RsbRB, RsbRC, RsbRD and YtvA (20). They form a large complex with the kinase RsbT and its antagonist or inhibitor, RsbS (24). We tested individual *rsbR* mutants in the presence of hydrogen peroxide using logarithmic growing pre-adapted cells and saw that *rsbRA* was equally sensitive to hydrogen peroxide as a *sigB* delete as previously shown (Figure 3A). Strains lacking *rsbRB* that were preadapted with ethanol exposure were more sensitive to ROS lethal levels than wild type cells (ANOVA P value <0.0001 and Dunnett’s test showed statistical significance). RsbRB is a co-antagonist of RsbT activation, similar to RsbRA, and strains lacking *rsbRB* have elevated SigB-dependent expression in presence of ethanol exposure (20). This suggests that in our experiments, SigB activity is elevated, yet it was not sufficient to protect cells against ROS, therefore the proper modulation that the second co-antagonist, RsbRB, provides is important for surviving lethal oxidative stress. While RsbRB can be a co-antagonist (21, 34), it may need other paralogs for proper regulation as our assay shows that RsbRB function is necessary for survival even when other co-antagonists are present. Deletion of *rsbRD* also made cells sensitive to lethal ROS even in the presence of ethanol preadaptation (Figure 3A). Interestingly, cells lacking *rsbRD* have no reported defect in SigB activation (20), yet there was a statistically significant difference between *rsbRD* null and wild type cells (ANOVA P value <0.0001, Tukey’s and Dunnett’s test showed statistical significance). While we do not know how the lack of *rsbRB* affects the stressosome, our results suggest that its presence in the complex plays a role in the regulation of SigB activity during ROS cross-protection.

**Figure 3.**
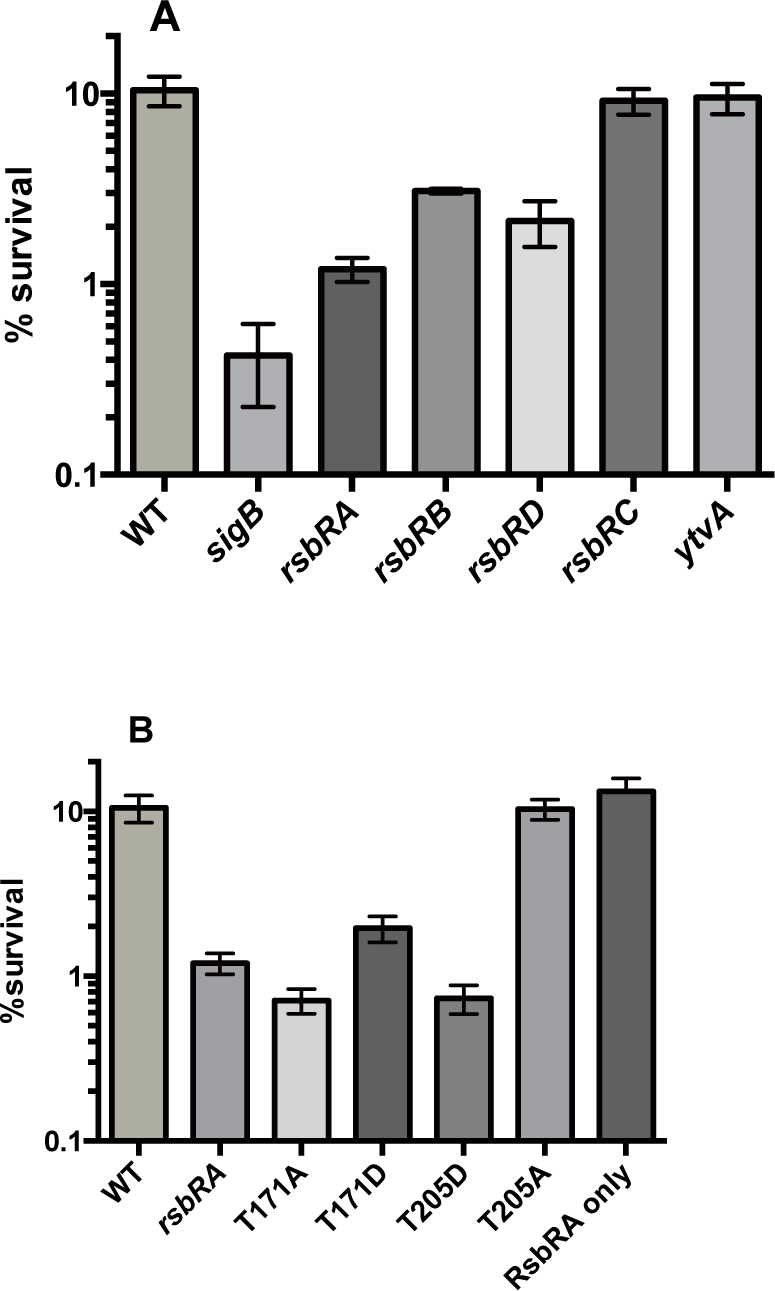
RsbRA phosphorylation is required for protection against oxidative stress. (A) Bacterial survival was measured in BLM, after 2% ethanol for 20 minutes and subsequent 5 mM hydrogen peroxide for another hour. Cells were allowed to recover overnight on LB plates. All mutants were tested and compared to wild type and *sigB.* The percent survival for each mutant was calculated compared to wild type survival (100%): *sigB* 5%, *rsbRA* 9%, *rsbRB* 52%, *rsbRC* 75%, *rsbRD* 43% and *ytvA* 82%. (B) Position of single amino acid mutations are labeled, RsbRA only strain has deletions in *rsbRB, rsbRC* and *rsbRD.* Every experiment was done a minimum of three times and averaged; standard error bars are shown for all experiments. One-way ANOVAs were performed, followed by Tukey’s and Dunnett’s multiple comparison tests to determine statistical significances between means.

In contrast, deletion of *rsbRC* had no effect on the cells’ ability to be cross-protected against ROS, showing viability undistinguishable from wild type. (Figure 3A). This is consistent with the absence of a recorded phenotype for cells lacking *rsbRC* (20). Interestingly, cells that contain RsbRC as the only co-antagonist in the stressosome have elevated SigB expression (21, 34) arguing that RsbRC alone is defective at preventing RsbT activation. And in the case of our experiments, removing RsbRC from the stressosome had no effect on the physiological outcome of stress cross-protection. Therefore, RsbRC is not necessary for cross-protection likely due to the redundancy of the paralogs in the complex. Similarly, deletion of *ytvA* had no effect on stress induced, cross-protection against hydrogen peroxide (Figure 3A). YtvA plays a role in the ability of cells to detect light, and cells without *ytvA* have reduced SigB activation under normal laboratory lighting conditions (20). Since our experiments were performed under similar lighting conditions, *ytvA* nulls likely had compromised signaling, yet the predicted lower SigB activity did not prevent cross-protection.

### RsbRA phosphorylation is important in the survival to oxidative stress

Since we saw a defect in rsbRA-deleted cells’ ability to cross-protect, we tested whether the known phosphorylation steps were involved during ROS exposure. First, we saw that cells where the stressosome consisted of only RsbRA were fully capable of surviving oxidative stress (Figure 3B) suggesting that at least during oxidative stress survival, the other RsbR proteins are not necessary and signaling through RsbRA is sufficient. Mutations in RsbRA phosphorylation site T171, T171A and T171D, made cells deficient at stress induced, ROS protection in our assay (Figure 3B). T171A and T171D mutants are known for having significantly diminished SigB activation measured by *ctc* expression (21). Our sensitivity results are consistent with these mutants having compromised SigB activation when cells were treated with ethanol, which resulted in lower SigB dependent expression of important genes, making cells sensitive to subsequent ROS treatment. Moreover, T171D mutant cells have lower SigB activity in the presence of salt stress compared to wild type, and the T171A mutant RsbRA protein was unable to promote RsbS phosphorylation by RsbT *in vitro* (27, 35), suggesting that the low *ctc-lacZ* expression in these mutants could have been due to lack of RsbS phosphorylation and failure to activate the stressosome or RsbT. These results are consistent with our cross-protection data showing mutations in T171 made cells sensitive to oxidative stress likely due to defects in stressosome priming and eventual cross-protection.

Mutations in the phosphorylation site T205 to Alanine or Aspartic acid had different phenotypes likely due to the previously observed effects of each amino acid substitution. First, the T205A mutation had no observable effect in our stress induced, ROS protection survival assay (figure 3B). T205A mutant cells were shown to have wild type levels of SigB dependent expression under 4% ethanol (21), which is higher than the priming stress we used, 2% ethanol. Therefore, SigB activation is likely normal in the T205A mutant and cells had sufficient SigB activity to protect them against subsequent lethal oxidative stress. On the other hand, the T205D mutant was very sensitive to oxidative stress cross-protection (Figure 3B) showing sensitivity similar to *rsbRA* null cells and is consistent with the effect of this mutation on SigB dependent expression since T205D mutant cells have lower SigB dependent expression than wild type cells in presence of salt and ethanol stress (21,27, 36). It is likely that in our viability assay, 2% ethanol did not cause SigB activation in this mutant therefore, ROS cross-protection could not happen and cells became as sensitive as *rsbRA* as Figure 3B demonstrates.

### RsbT and RsbP are important during disulfide stress cross-protection

Disulfide stress happens when thiol groups on proteins are oxidized and non-native covalent bonds form disrupting protein function. Spx and MgsR are disulfide stress regulators responsible for regulation of multiple genes involved in the detoxification of disulfide stress (37, 38). SigB controls their induction during ethanol stress therefore, environmental stress priming could also protect against disulfide stress. Using diamide to induce disulfide stress, *sigB, rsbT* and *rsbP* null cells were tested in cross-protection during disulfide stress in logarithmically growing cells. We saw that *sigB* deleted cells were defective in survival during diamide exposure compared to wild type cells and preadaptation heightened this difference between wild type and *sigB* null cells (Figure 4A). Similarly, *rsbT* mutants showed lower survival than wild type, whereas *rsbP* mutant cells survived to wild type levels (Figure 4B). In stationary phase, which induces nutritional stress, *sigB* and *rsbP* deleted cells were more sensitive to diamide exposure than wild type and *rsbT* deleted cells (Figure 4C). Therefore, nutritional and environmental stress prime cells against disulfide stress.

**Figure 4.**
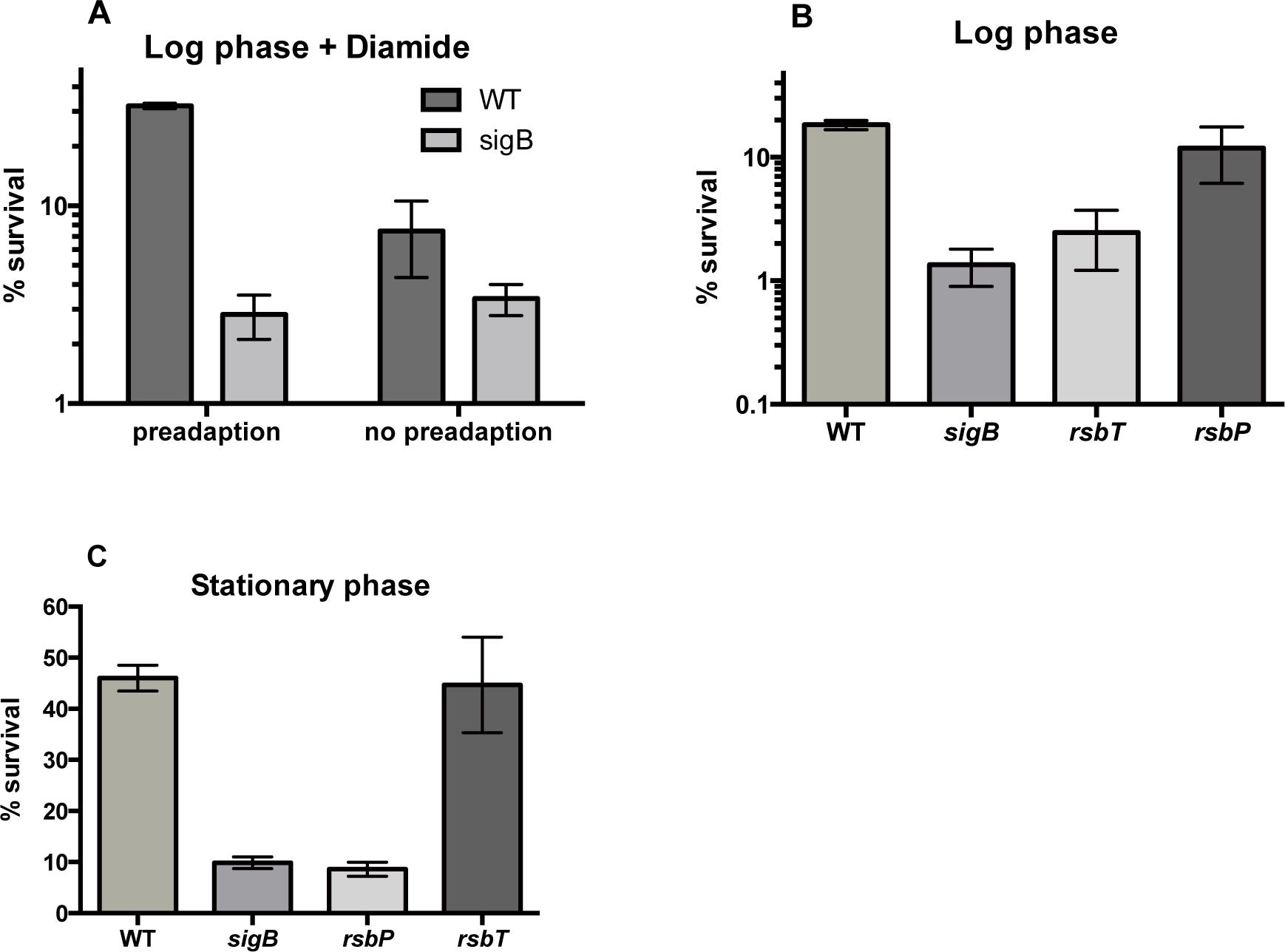
Environmental and nutritional stress protects against disulfide stress. (A) Cells were grown in BLM until OD600nm 0.4 and were either preadapted with 2% ethanol for 20 minutes or received no preadaptation. Diamide was added at 45 mM final concentration for 1 hour before cells were allowed to recover overnight on LB plates. (B) Cells were grown to mid-log in BLM, treated with 2% ethanol for 20 minutes and then given 45 mM diamide for one hour. (C) Cells were grown in Glucose Limitation Media and monitored until transition state. One hour into transition state, cell were given 45 mM diamide for one hour. Every experiment was done a minimum of three times and averaged; standard error bars are shown for all experiments. One-way ANOVAs were performed, followed by Tukey’s and Dunnett’s multiple comparison tests to determine statistical significant differences between means.

### Role of SigB in resilience to nitrosative stress

We tested how general the oxidative stress cross-protection imparted by SigB was by exposing cells to nitrosative stress. Viability after SigB activation was measured by treating cells with the NO producing compound Sodium Nitropruside (SNP) during log phase or during early stationary state to measure the role of each SigB activating pathway. Wild type and sigB-deleted cells were pretreated with ethanol to activate the stressosome and then SNP was added for one hour. As shown in Figure 5A, pre-treatment in log phase made the cells more resistant to lethal levels of SNP, and this resistance was SigB-dependent. SigB-dependent survival to SNP was not observed in previous experiments by Rogstam et al. (12) but the growth medium and stress conditions used in their study and ours were significantly different. We use Basic Limitation Medium and they used Nutrient Sporulation Medium. Additionally, the adaptive response they tested used low level exposure to 0.5 mM SNP followed by lethal SNP levels, whereas our assay uses an unrelated stressor, ethanol, to activate the environmental stress priming effect. Therefore, under their conditions the general stress response was potentially not activated compared to SigB activation in our system using ethanol.

**Figure 5.**
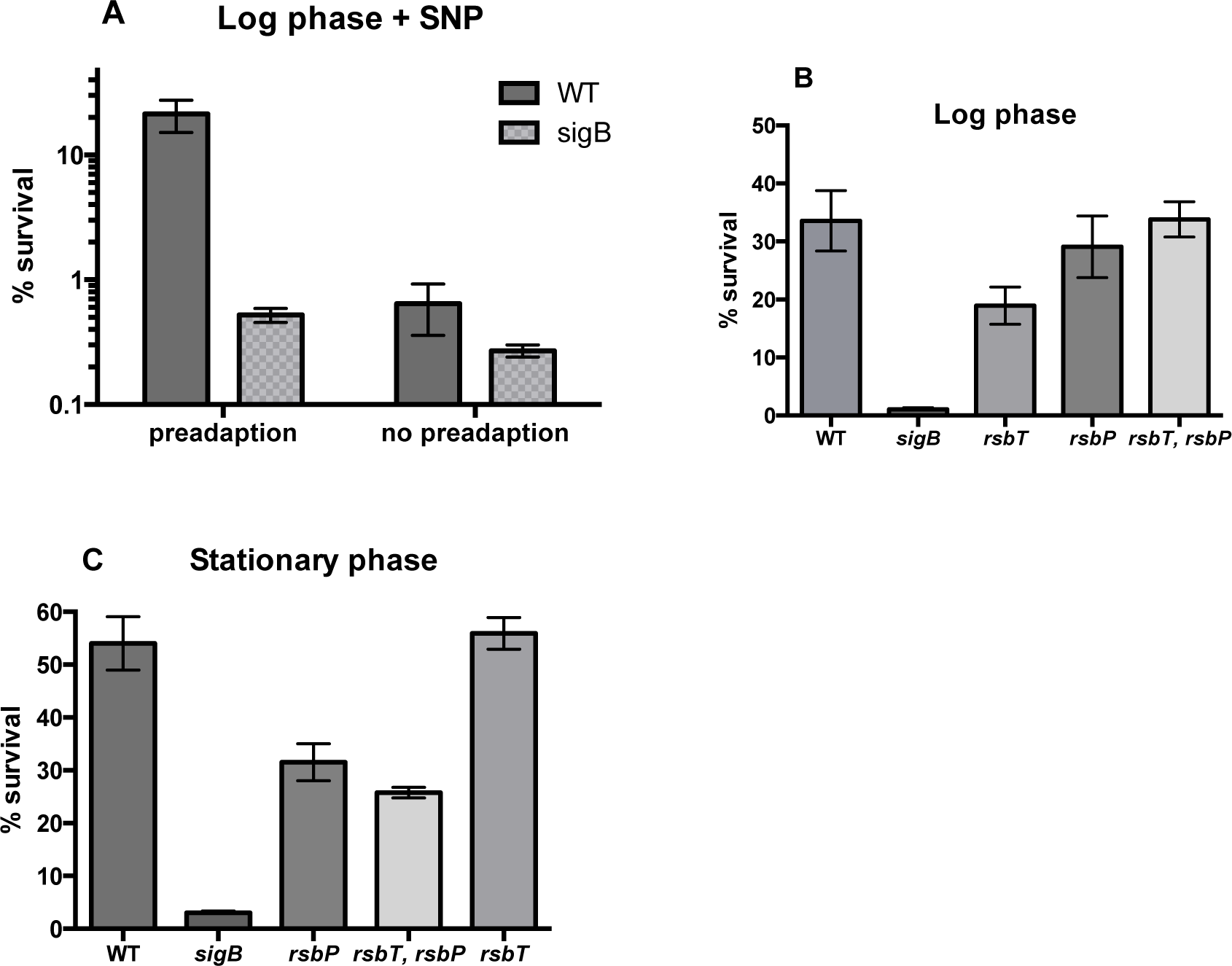
Environmental and nutritional stress protects against nitrosative stress. (A) Cells were grown in BLM until midlog and were either preadapted with 2% ethanol for 20 minutes or received no preadaptation. Sodium nitropruside was added at 74 mM final concentration for one hour before being plated on LB overnight. (B) Cells were grown to midlog in BLM, treated with 2% ethanol for 20 minutes and then given 74 mM sodium nitropruside for one hour. (C) Cells were grown in Glucose Limitation Media and monitored until transition state. One hour into transition state, cell were given 74 mM sodium nitropruside for one hour and then plated. Every experiment was done a minimum of three times and averaged; standard error bars are shown. One-way ANOVAs were performed, followed by Tukey’s multiple comparison tests to determine statistical significant differences between means.

To test the upstream activators of the priming pathway, mutants in *rsbT* and *rsbP* were subjected to the cross-protection assay. Wild type and rsbP-deleted cells had similar viability rates supporting that RsbP is not activated in log phase by ethanol stress (Figure 5B). Interestingly, cells with deleted *rsbT* showed an intermediate phenotype between wild type and *sigB* mutants (ANOVA P value 0.0001. Tukey’s test found no difference between wild type and *rsbT* cells but a significant difference with *sigB* cells). It is possible that under conditions where the stressosome is non-functional, such as in the *rsbT* deleted cells, RsbP becomes required. To test this hypothesis we treated *rsbT, rsbP* double mutant cells with SNP and saw that this strain had similar viability to the rsbP-deleted cells, and not like cells lacking *sigB* as was expected if our hypothesis was correct (Figure 5B). Instead, this result suggests a SigB-dependent cross-protection pathway that does not use the known activators RsbT and RsbP. While we have not tested the genetic requirement of downstream regulators such as RsbV in our experiments, the data suggest that SNP causes damage that can be survived if SigB is activated by environmental stress, suggesting an alternative pathway to activate SigB in log phase. RsbV-independent activation was observed during chill (15°C) and high temperature (51°C) exposure (39, 40). In these temperatures, *rsbV-deleted* cells had higher than usual SigB protein levels as if deleting these regulators causes hyper-activation of SigB, which would also explain our viability results in the double mutant strain. Whether there is another branch of the general stress signaling network is worthy of further investigation.

In stationary phase, cells were treated with lethal levels of SNP and viability was measured. Wild type and *rsbT-deleted* cells showed the same resistance as with other types of oxidative stress arguing that in stationary phase the stressosome is not required (Figure 5C). The single *rsbP* mutant and the double *rsbP, rsbT* mutant were less sensitive than *sigB* deleted strains when exposed to lethal SNP concentrations (Figure 5C). Both results suggest that SNP resistance may require SigB activation that happens through a pathway other than the known RsbV anti, anti-sigma factor, since so far only the phosphatase activity of RsbP and the stressosome-activated RsbU are required for RsbV activation. Alternatively, SNP may cause RsbV activation through a yet uncharacterized mechanism, which works in both logarithmic and transition state. We have shown that SNP causes stress that requires SigB activity for optimal survival but the mechanism of SigB activation under nitrosative stress remains unknown.

## Discussion

The general stress response activated by SigB gives cells an advantage to uncertain, future environmental conditions. We characterized the SigB regulatory pathways required for enhanced survival during oxidative stress due to environmental and nutritional stress priming. We showed that upstream regulators of SigB are involved in *B. subtilis* stress priming against oxidative stress, disulfide stress and reactive nitrogen species and provide evidence that stressosome components, RsbRB and RsbRD, may play a role in ROS signaling outside of environmental stress SigB activation. Bacteria have multiple strategies to deal with their natural ecosystems, these include slowing down metabolism during transition state, inducing competence, biofilm formation, sporulation and virulence in pathogenic bacteria. Since SigB affects some of these processes (41, 42) it is possible that priming is also involved in these distinct states. If low-level SigB activity gives cells an advantage, then normal environmental fluctuations in temperature, osmotic pressure and carbon limitation might help cells more successfully transition between developmental and life style states. Moreover, endogenously produced radicals through metabolic reactions and aerobic respiration must be detoxified (43) and SigB could play a more important role in ROS and RNS detoxification than previously thought. In pathogens redox sensing of the extracellular environment is essential to survival, and for those species that express SigB, it appears to be important in the initial steps that lead to successful colonization (1, 44). In their natural environments, populations may experience sporadic SigB activation due to small changes in temperature or pH and these changes may prepare the cells for extreme oxidative conditions such as the ones imposed by the immune system.

### Role of the stressosome in modulating SigB activity during oxidative stress

We saw that deregulated SigB-dependent transcription was counter-productive to the benefits of priming. Using viability as a measure for proper SigB function, we were able to separate mutations in stressosome genes into three categories. Mutations compromised at the priming step were most sensitive, *rsbRA* null, *rsbRA* T171A, T171D and T205A, and had viability similar to *sigB* nulls, as expected if their only role was in priming. Mutations that were priming-proficient but oxidative stress sensitive, such as *rsbRB* and *rsbRD* suggest a priming-independent role in ROS sensing or signaling for the stressosome that has never been observed. Finally, mutations in *rsbRC* that retained the ability of cells to be primed even to a lower degree, as in *ytvA* nulls, survived oxidative stress like wild type cells. The redundancy of stressosome proteins could be at play during priming so that *rsbRC* and *ytvA* null cells activated SigB to sufficient levels.

Using an assay that measures the physiological effects of oxidative stress exposure, we were able to show a novel phenotype for two stressosome genes, *rsbRB* and *rsbRD* that cannot be explained by a lack of priming. Cells with mutations in *rsbRA* that reduced SigB activity, were less efficient at oxidative stress cross-protection (Figure 3B) as expected if priming is the only role the stressosome plays. Yet, mutations that induce SigB activity such as deletion of the stressosome antagonist protein RsbRB lowered the cell’s resilience or ability to meet subsequent oxidative stress. We propose two alternative explanations for this observation. First, hyper-active SigB signaling could be detrimental to the expression patterns required for cross-protection by some general disruptive mechanism of imbalanced gene products. Alternatively, RsbRB and/or RsbRD proteins could have a direct or indirect role in sensing oxidative stress, which contributes to the cross-protection we observed. While, no sensing mechanism has been described for the *B. subtilis* stressosome, both direct and indirect sensing functions have been reported in *Vibrio brasiliensis* (45) and *Listeria monocytogenes* (46) stressosomes. In the *Vibrio* system, the RsbR co-antagonist bound oxygen, which could make this species stressosome an oxidative stress sensing complex (45). *L. monocytogenes* stressosomes did not directly bind a ligand, but a transmembrane protein, Prli42, directly interacted with RsbRA and was required for SigB dependent expression during hydrogen peroxide exposure (46). This mechanism could be conserved in *B. subtilis,* making RsbRB and RsbRD interesting candidates for oxidative stress signal transducers.

### Cross-protection and SigB regulatory pathways

SigB’s importance in oxidative stress cross-protection was first appreciated for its contribution to transition state (15) and later for its role in logarithmic growth (10). While oxidative stress resistance is known to be SigB dependent, we provide evidence that in stationary phase RsbP is the most important SigB regulator for priming and RsbT plays a more significant role in logarithmic phase. During nutritional stress the potential redox imbalance caused by depletion of ATP could be sensed and processed by the two functions in the RsbP-RsbQ complex. The PAS domain on RsbP could bind the signal molecule (47) and RsbQ’s hydrolase domain could process it; yet imbalanced redox state was not involved in the activation of RsbP arguing against the redox sensing model (19). However, our cross-protection experiments revealed a potentially uncharacterized SigB activating pathway involved in oxidative stress caused by reactive nitrogen species (Figure 5). Nitrosative stress is an inducer of SigB-dependent gene expression (12, 48, 49). In aerobic conditions, *rsbT* and *rsbP* were each required depending on mode of NO production (49) so how the stress signal(s) activates SigB remains unknown. Our results are consistent with this observation because we saw a decrease in survival in rsbT-deleted cells, although not to the extent of *sigB* deleted cells. Moore and colleagues measured SigB-dependent transcription, so a direct comparison is difficult given that our assay measures the physiological effect of SigB activation. Importantly, we saw that nitrosative stress cross-protection required SigB but not necessarily RsbT or RsbP (Figure 5) arguing for an RsbV-independent pathway or regulation of RsbV independent of the known phosphatases. It is known, however that chill and high temperature induce SigB in an RsbV-independent way (39, 40). While we do not know whether nitrosative stress activates SigB through the same pathway used by extreme temperatures, these results together raise the possibilities that SigB can be activated by more uncharacterized mechanisms.

### General Stress Response and Antioxidant Activity

Disulfide stress sensing is conserved in many bacterial species through the disulfide sensing, transcription factor Spx. It is responsible for regulating genes such as thioredoxins that reduce inappropriate disulfide bonds between proteins (37). Since Spx is under the regulation of SigB during ethanol stress (50), its activation could explain the cross-protection, i. e. resilience, observed when cells were treated with lethal amounts of diamide (Figure 4). Likewise, the Spx homolog, MgsR is regulated transcriptionally by SigB (38). The sensitivity of sigB-deleted cells to diamide exposure could be explained if transcription factors, Spx and MgsR, were not induced. Additionally, some MgsR regulated genes have SigB dependent promoters (38), making their transcription both directly and indirectly sensitive to SigB activity. Appropriate Spx and MgsR activity levels could be required for the concerted transcription of SigB-dependent genes with potential detoxification properties such as predicted dehydrogenases and reductases regulated by Spx and MgsR (38).

Nitric oxide production by SNP and diamide stress cause disulfide bond intermediates (51) that result in non-native disulfide bonds requiring detoxification and antioxidant activity for survival. *B. subtilis* produces bacillithiol, the low molecular-weight thiol, involved in redox chemistry. It is synthesized by acillithiol biosynthethic enzymes and transferred to toxic substrates for detoxification by Bacillithiol-S-Transfereses (52, 53). Two bacillithiol transferase genes, *bstB* and *bstD* show mRNA expression patterns similar to SigB-dependent genes, high in ethanol, heat, hydrogen peroxide and diamide exposure (54) yet they are not known SigB-targets. If *bstB* and *bstD* expression is induced by environmental stress conditions, they could be indirect targets of SigB through MgsR activity, contributing to the SigB dependent survival we observed during priming. Consistent with a detoxifying role of bacillithiol in disulfide stress, the promoters of bacilithiol biosynthetic genes, *bshA, bshB1/2, BshC,* are upregulated by Spx during difulside stress (53). Ultimately, stress priming triggered through ethanol exposure could induce bacillithiol synthesis and utilization promoting the enhanced resistance of cells subsequently exposed to toxic diamide and nitrosative stress. Potentially, SigB regulatory proteins such as the stressosome and the RsbP/RsbQ complex could function in the cross-protection to all types of oxidative stress conditions, providing primed antioxidant capabilities to the cell.

## Acknowledgements

This research was supported by the M.J. Murdock Natural Science Grant #2014262 awarded to Carla Y. Bonilla and Howard Hughes Medical Institute through the Undergraduate Science Education Program awarded to Gonzaga University. The authors would like to thank Dr. Chester Price for generously providing *rsbRA* mutant strains and Dr. Alan Grossman for *Bacillus subtilis* strain and pGK67 plasmid. The authors would also like to thank Dr. Antonio Abeyta and Dr. Kirk Anders for comments on this project and Dr. Price for critical reading and feedback on this manuscript.

